# Lecanemab binds to transgenic mouse model-derived amyloid-β fibril structures resembling Alzheimer’s disease type-I, type-II and Arctic folds

**DOI:** 10.1101/2025.01.17.633637

**Authors:** Fernanda S. Peralta Reyes, Dieter Willbold, Gunnar F. Schröder, Lothar Gremer

## Abstract

Lecanemab, an Alzheimer’s disease US Food and Drug Administration approved monoclonal antibody was previously reported to have a high affinity against intermediately sized amyloid-β aggregates. Subsequently, it was observed by immunogold labelling that lecanemab can also bind to human type-I amyloid-β fibrils. Therefore, to determine whether lecanemab binds to amyloid-β fibril structures other than type-I, we performed immunogold labelling on extracted amyloid-β fibril preparations from six different Alzheimer’s disease mouse models whose structures were previously solved by cryo-EM. Our results show that lecanemab exhibits high binding affinity to amyloid-β fibril structures that have a flexible N-terminus in common, as it is the case for type-I, type-II and murine type-III amyloid-β fibril polymorphs which resemble or are identical to human structures observed in sporadic and familial cases of Alzheimer’s disease, including a case with the Arctic (E22G) mutation. In contrast, only weak, if any, lecanemab binding was observed for amyloid-β fibril folds with a fixed and ordered N-terminus.

**Key points:**

- Lecanemab binds to Aβ fibrils from several Alzheimer’s disease tg-mice whose structures resemble the type-I, type-II and Arctic folds found in Alzheimer’s patients, all of which share a flexible, unstructured N-terminus.
- Lecanemab is therefore expected to be active against all common familial and sporadic Alzheimer’s cases containing these folds.
- Lecanemab binding ability is unaffected by and tolerates the Arctic E22G mutation, at least in type-I or Arctic folds.
- Weak, if any, lecanemab binding was observed to Aβ fibrils derived from tg-SwDI mice, whose structures DI1, DI2 and DI3 all share structured and fixed N-termini.
- Since the fixed N-termini of tg-SwDI DI1 fibrils and human meningeal Aβ40 fibrils derived from CAA-affected brain are identical, most likely preventing lecanemab binding, treatment with lecanemab may be less or ineffective against CAA, but may explain the reported beneficial low ARIA-E frequency with this antibody.

**Table of Contents Graphic (TOC):** 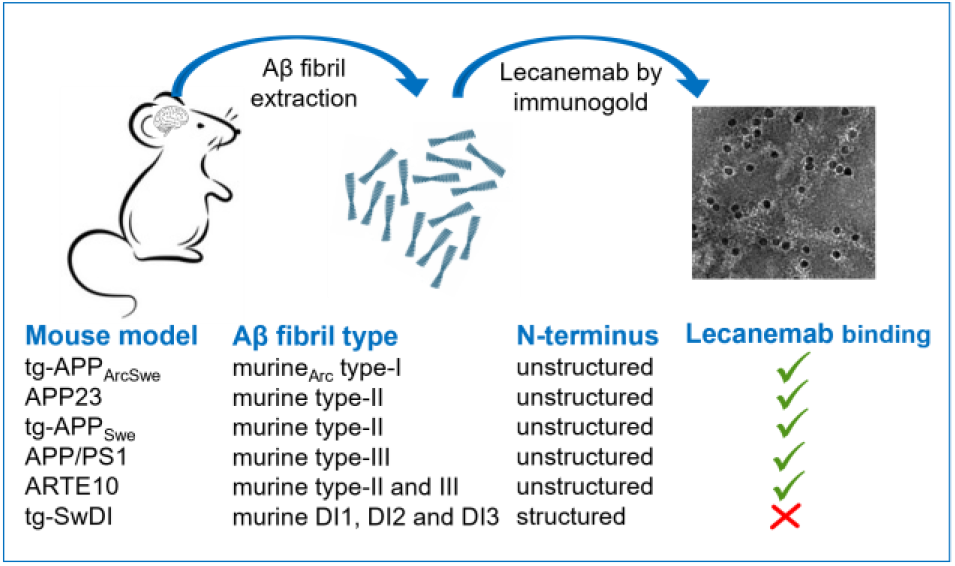

## Introduction

Alzheimer’s disease (AD) is the most common type of dementia and is pathologically associated with the presence of extracellular amyloid-β (Aβ) plaques and intracellular neurofibrillary tangles. When the amyloidogenic pathway is carried out, it results in the formation of monomeric Aβ which is the building unit for the assembly of oligomers, protofibrils, fibrils and plaques^1^. Different therapeutic options intend to target different Aβ species, however, many of them have failed to show clinical efficacy, while others, such as donanemab and lecanemab (a.k.a. BAN2401) are fully approved by the US Food and Drug Administration (FDA)^2,3^. Additionally, aducanumab has also received an accelerated conditional FDA approval^3^.

mAb158, the murine predecessor antibody of lecanemab, was mainly designed to bind soluble protofibrils rather than mature and insoluble fibrils. However, ELISA experiments showed that mAb158 also binds Aβ fibrils, suggesting that the epitope present in protofibrils is also present in fibrillar structures. Nevertheless, mAb158 has no affinity for the Aβ protein precursor (AβPP) and did not bind fibrils from other amyloids^4,5^.

Furthermore, it was reported that the humanized IgG1 antibody lecanemab can bind with high affinity to soluble protofibrils and only with moderate selectivity to Aβ fibrils when compared to monomeric Aβ^6^. Soluble protofibrils were portrayed in more than one occasion as the most toxic Aβ species^7^, making lecanemab a high profile therapeutic option. During a phase-three clinical trial, lecanemab administration to patients in the early stage of the disease showed decreased amyloid levels in the brain and a moderate reduction of cognitive decline when compared to placebo^6^. When interpreting the data regarding amyloid levels in the brain, it is worth to highlight that lecanemab does not interfere with the Positron Emission Tomography (PET) radioligand [^11^C]-Pittsburgh compound B when binding to Aβ deposits, as both have different binding sites^8^.

Although it was reported that lecanemab does not have high binding affinity to fibrils^6^, it was shown by immunohistochemistry that lecanemab also stains Aβ plaques^9^. Additionally, it was portrayed by immunogold-electron microscopy (immunogold-EM) that lecanemab can indeed bind Aβ fibrils that were observed in ultracentrifugal supernatants of aqueous extracts from the human brain parenchyma^10^ as shown for two samples containing mainly Aβ fibrils denominated as type-I^10^, observed in sporadic and familial AD cases^11,12^. Whether lecanemab can also bind to other pathologically relevant types of Aβ polymorphs remained to be elucidated.

Therefore, we analysed lecanemab’s binding competence by immunogold-EM on various *ex-vivo* Aβ fibril polymorphs. These Aβ fibrils were derived from brain samples of six common AD tg-mouse models in the pre-clinical context, which structures have been recently solved by cryo-EM^13^. Our selection include the tg-APP_ArcSwe_ tg-mouse model which was efficiently used in the pre-clinical evaluation of lecanemab^14^ and is so far the only model whose Aβ fibrils resemble the human type-I Aβ polymorph. Additionally, AD tg-mouse models that exhibit fibrils of the human type-II polymorph, mainly observed in familial AD cases and other conditions, as well as tg-mouse models resembling the human Arctic fibril fold, and finally a tg-mouse model with other novel Aβ structures were assessed^13^.

## Animals and ethical approval

The *ex-vivo* Aβ fibril sample preparations analysed in the present study were previously used to solve their cryo-EM structures^13^ and were isolated from the following mouse models: APP/PS1 (APPswe/PSEN1dE) (heterozygous; *n* = 1 (male); 33 months old) on a C57BL/6;C3H background. ARTE10 (homozygous; *n* = 1 (female); 24 months old) on a C57Bl/6 background which was a gift from Taconic Biosciences. Tg-SwDI (heterozygous; *n* = 1 (male); 29 months old) on a C57BL/6 background which is not only used as a model for AD but also for cerebral amyloid angiopathy (CAA). APP23 (heterozygous; n = 1 (male); 21 months old) on a C56BL/6 background. Tg-APP_ArcSwe_ (heterozygous; n = 1 (male); 18 months old) and tg-APP_Swe_ (heterozygous; n = 1 (male); 22 months old), both on a C57BL/6 background.

Experiments that were performed on the APP/PS1, ARTE10, tg-SwDI and APP23 AD mouse models were conducted in line with the German Law on the protection of animals (TierSchG §§7–9). APP/PS1 mice breeding was validated by a local ethics committee (Landesamt für Natur, Umwelt und Verbraucherschutz Nordrhein-Westfalen (LANUV), Az: 84-02.04.2019.A304). APP/PS1 and tg-SwDI mouse lines were acquired from the Jackson Lab (JAX MMRRC Stock no. 034829 or JAX MMRRC Stock no. 034843). Breeding of tg-APP_ArcSwe_ and tg-APP_Swe_ mouse models was done under the ethical permit 5.8.18-20401/20, approved by the Uppsala County Animal Ethics Board. All mice were treated under controlled conditions at 22 °C, 12:12 h light: dark cycle, 54% humidity, as well as food and water *ad libitum*.

## Materials and Methods

Aβ fibril extraction was done based on a well-stablished sarkosyl-based approach^12,13^. Between 0.4 and 0.6 g of non-fixed brain tissue from six different AD tg-mouse models was snap-frozen in −80°C cold isopentane and stored at −80°C. The tissue was then thawed and physically homogenized in a 20-fold volume (w/v) of extraction buffer (10 mM Tris-HCl, pH 7.5, 0.8 M NaCl, 10% sucrose, 1 mM EGTA) using a Dounce glass tissue grinder. 10% aqueous sarkosyl (Sigma-Aldrich) was added to bring the brain homogenate to a final sarkosyl concentration of 2%. The sample was mixed thoroughly by pipetting up and down 30-times before incubation at 37°C for 1 h. The homogenate was then centrifuged at 10,000x*g* in a table-top centrifuge at 4°C for 10 min. The pellet was discarded and the supernatant was ultracentrifuged at 100,000x*g* at 4°C for 1 h (Beckman Coulter Optima MAX-XP, TLA55 fixed-angle rotor). The resulting supernatant was discarded and the pellet was resuspended and mixed with extraction buffer (1 ml g^−1^ original tissue mass) before low-speed centrifugation at 5,000x*g* at 4°C for 5 min. Afterwards, the supernatant was threefold diluted in dilution buffer (50 mM Tris-HCl, pH 7.5, 0.15 M NaCl, 10% sucrose, 0.2% sarkosyl) and ultracentrifuged once more at 100,000x*g* at 4°C for 30 min. The final Aβ fibril-rich pellet was resuspended (100 µl g^−1^ original tissue mass) in resuspension buffer (20 mM Tris-HCl, pH 7.4, 50 mM NaCl), frozen in liquid nitrogen and stored at −80°C for further use.

Immunogold labelling was performed according to the protocol of Gulati et al^15^. In brief, 300-mesh carbon-coated copper grids (EM Sciences, ECF300-CU) were glow discharged with a PELCO easiGlow Glow Discharge Cleaning System. 3 µl sample of extracted Aβ fibril suspension were incubated for 2 min on the grid’s surface and excess liquid was blotted with filter paper afterwards. The grid was then placed on top of a 15 µl _d_H_2_O droplet on parafilm for 1 min and blotted. Afterwards, the grid was transferred to a 15 µl droplet of blocking buffer (99 ml PBS, pH 7.4, 100 μl Tween-20, 1 ml 30% IgG-free bovine serum albumin) inside of a humidifying chamber and incubated for 15 min. After blotting, the grid was transferred to a 15 µl droplet of lecanemab primary antibody diluted in blocking buffer to 2 µg ml^-1^ for 1-2 hours and blotted once more. The grid was washed five-times by incubating it in 15 µl droplets of washing buffer (100 ml PBS, pH 7.4, 100 μl Tween-20, 100 μl 30% IgG-free bovine serum albumin) for 3 min and blotting with filter paper after each wash. The grid was then transferred to a 10 nm gold-conjugated goat anti-human secondary antibody (Abcam) diluted 1:20 in blocking buffer droplet for one hour. The grid was washed five-times with washing buffer and three-times with _d_H_2_O as described above. The grid was then transferred to a 15 µl droplet of 1 % uranyl acetate for 1 min, blotted and air-dried before transmission electron microscope (TEM) imaging (ThermoFisher Scientific Talos-L120C).

## Results and Discussion

First, we tested by EM-immune-gold labelling whether lecanemab is able to bind *ex-vivo* murine type-I Aβ fibrils obtained from tg-APP_ArcSwe_ mouse brain tissue. The fibrils were incubated with lecanemab as the primary antibody, then reacted with the secondary antibody with conjugated 10 nm gold-nanoparticles, followed by negative-staining with uranyl acetate. The TEM electron micrographs reveal a specific fibril decoration with the electron-dense 10 nm gold-nanoparticles (Fig. 1A) indicating specific lecanemab binding to type-I Aβ fibrils from tg-APP_ArcSwe_ mouse brain. Remarkably, the tg-APP_ArcSwe_ mouse model was used for the pre-clinical validation of lecanemab^14^ and is the only tg-AD mouse model up to date that resembles the human type-I Aβ fibril polymorph^13^, present in sporadic^12^ and familial AD cases^11^ including Down syndrome^16^. Although the murine and human type-I Aβ fibrils show subtle differences due to the Arctic (E22G) mutation in the tg-APP_ArcSwe_ mice, their overall fold, side-chain orientation and fibril surface is highly conserved. And indeed, previous findings have shown that lecanemab binds to human *ex-vivo* type-I Aβ fibrils^10^ that do not carry any mutation, indicating that the presence or absence of the E22G mutation does not affect lecanemab binding.

**Figure 1.**
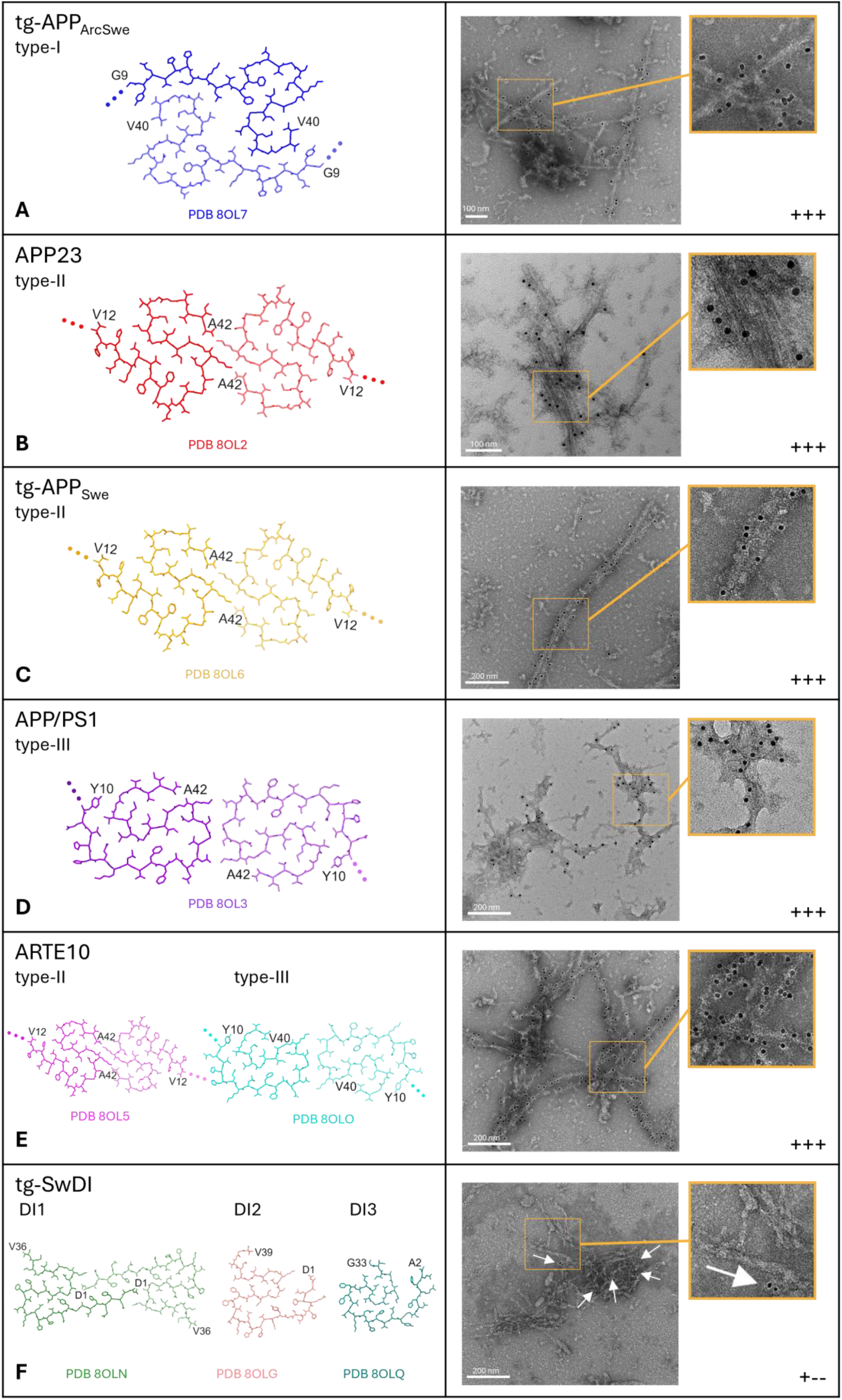
Lecanemab immunogold-labelling of Aβ fibrils with different molecular structures. Previously solved cryo-EM structures of Aβ fibrils from different AD mouse models^13^ (left). Immunogold TEM images of extracted Aβ fibrils with lecanemab as primary antibody (right). Approximate estimation of the amount of gold particles bound to the fibrils, where ‘+++’ means that gold-particles decorated the Aβ fibrils entirely, and ‘+--’ means that only few gold particles (white arrows) labelled the Aβ fibrils (bottom right). The dots indicate the unstructured, flexible N-terminal regions.

In familial AD and other conditions, another Aβ fibril fold, named type-II was identified as well^12,16^. Therefore, we were interested whether lecanemab can also bind and detect this fibril morph by using type-II Aβ fibrils extracted from brains from tg-APP_Swe_ or APP23 tg-mice, respectively^13^. After performing immunogold-EM, a clear gold labelling of the Aβ fibrils from both mouse models was observed, indicating high affinity binding of lecanemab to type-II fibrils (Fig. 1B,C). Considering that the 3D structures of type-II Aβ fibrils in humans and mice are identical down to atomic details^13^, we expect lecanemab to be effective against patients with AD pathologies involving Aβ type-II fibrils as well.

Furthermore, another Aβ fibril fold referred to as murine type-III was observed in Aβ fibril preparations from brain tissue of APP/PS1 and ARTE10 tg-mice, in the latter together with Aβ type-II fibrils^13^. Notably, the murine Aβ type-III fibril structure with its nonmutated Aβ42 sequence is highly similar to a protofilament pair involving protofilaments A and B from a tetrameric human Arctic Aβ fibril fold, despite the Arctic Aβ E22G mutation in the patient^17^.

After immunogold staining, the electron micrographs evidently show that lecanemab also binds and recognizes type-III fibrils present in APP/PS1 tg-mice (Fig. 1D). In addition, fibril preparations derived from ARTE10 tg-mice, which displayed both, type-II and type-III fibrils (Fig. 1E) also show high-affinity lecanemab binding, as expected.

Taking the similarity of the murine Aβ type-III fold and the human Arctic fold into account, our results may infer that lecanemab should also be effective in AD patients displaying this fold either in the nonmutated state (as in the APP/PS1 tg-mice), or in the E22G mutated state as in Arctic AD patients.

Other novel Aβ polymorphs denominated as DI1, DI2 and DI3, with DI1 as the most abundant were observed in Aβ fibril preparations from brains of SwDI tg-mice^13^ which serves as a model for both, AD and CAA. In contrast to type-I, type-II and the murine Aβ type-III folds, which all have in common a flexible N-terminus, all three resolved SwDI folds exhibit well-ordered and fixed N-termini^13^ (Fig. 1F).

Analysis of lecanemab’s binding capability to SwDI tg-mice derived Aβ fibrils by immunogold-EM revealed only very few gold particles bound to SwDI fibrils (Fig. 1F), which indicates a weak or negligible lecanemab binding, if present at all.

Considering that lecanemab’s binding site is reported on the N-terminus (between residues 1-16)^18^, the results indicate that the N-terminus needs to be flexible and non-structured to act as an efficient lecanemab binding epitope, as it is evident from high affinity binding of lecanemab to the type-I, type-II and the murine Aβ type-III folds (Fig. 1A-E) and weak or nonbinding to the SwDI folds (Fig. 1F). Interestingly, when compared to immunogold labelling using NAB228 as primary antibody, which also binds to the N-terminus (1-11), a similar pattern was observed^13^.

Even though the tg-SwDI folds are unlikely to be present in humans due to the double, Dutch (E22Q) and Iowa mutations (D23N) ^11^, the N-termini of SwDI DI1 and Aβ40 fibrils extracted from the human meninges^13,19,20^ are structurally highly similar (Fig. 2) ^13^. Therefore, lecanemab may have less binding affinity to CAA cases, as it was also confirmed in a previous study by Söderberg and colleagues^21^, where it was also observed that lecanemab has a relatively low amyloid-related imaging abnormalities with edema (ARIA-E) frequency (12.6%) when compared to other antibodies such as aducanumab, bapineuzumab, donanemab and gantenerumab which rather have higher ARIA-E frequencies (25-35%) and higher binding affinity to CAA fibrils^21^.

**Figure 2.**
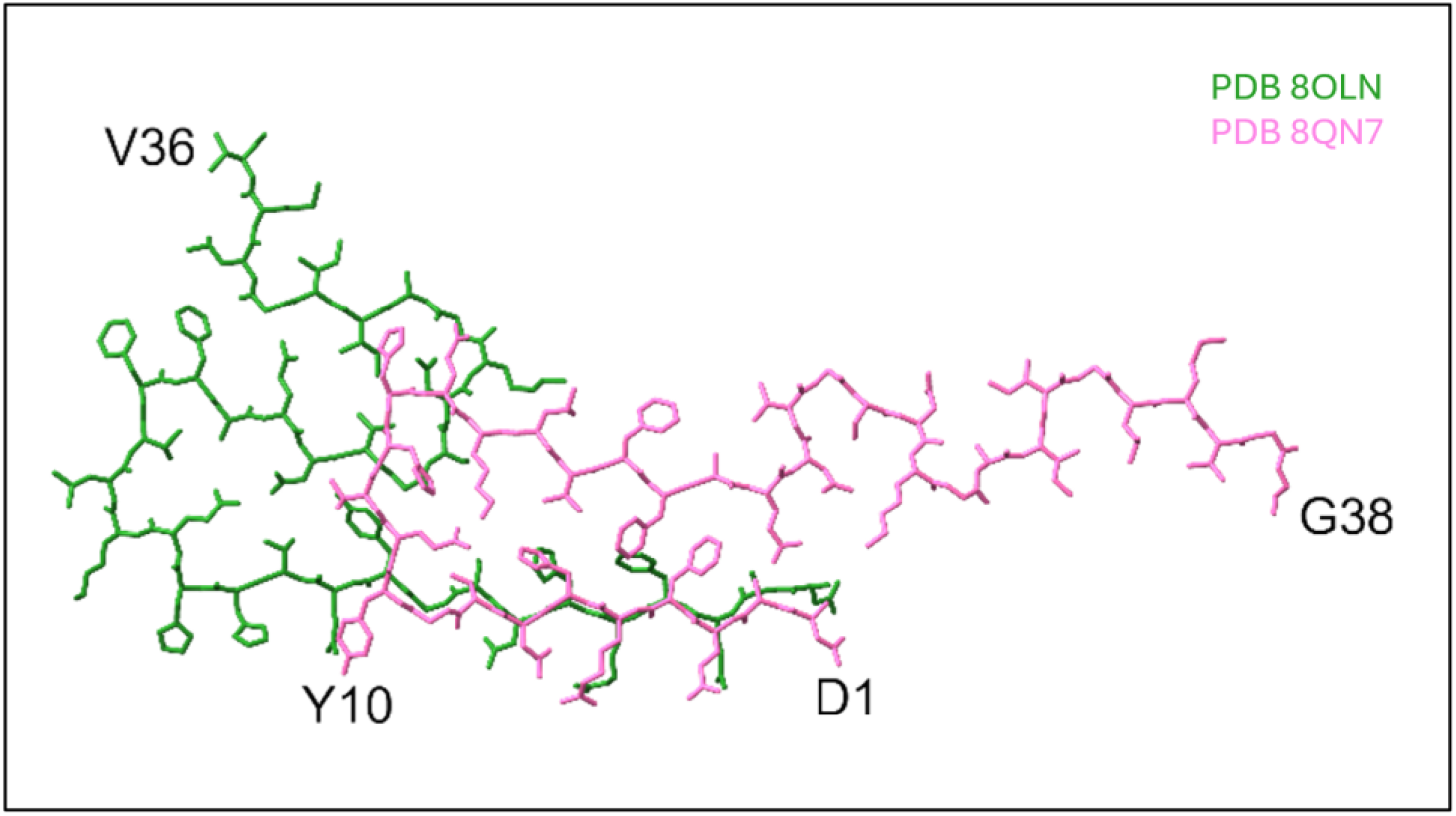
Structural comparison of DI1 Aβ fibrils with human CAA type Aβ40 fibrils. Overlay of the DI1 Aβ fibril structure from tg-SwDI mice (green; PDB 8OLN) with the cryo-EM structure of Aβ40 fibrils extracted from the leptomeninges of human brain tissue from a patient with Alzheimer’s disease (pink; PDB 8QN7) showing the similarity of the structured N-termini.

In conclusion, our results show that lecanemab is expected to be active against all common familial and sporadic AD cases containing type-I, type-II, or the Arctic fold, or mixtures of them, all having a flexible N-terminus in common (Fig. 1A-E). As lecanemab binds the type-I Aβ fibril fold in its nonmutated (human) as well in the E22G-mutated state (tg-APP_ArcSwe_ mice), it withstands the Arctic E22G mutation, and likely as well other E22 or neighbouring mutations (i.e. Flemish A21G, Dutch E22Q, Italian E22K, Iowa D23N), as long as the fibril fold is maintained. This principle may not be restricted to type-I fibrils, as the type-III fold (non-mutated in mice) and the human Arctic E22G fold are also structurally similar. Further research may focus on a detailed structural characterisation of the lecanemab binding modes to the various AD relevant Aβ fibril folds.

## Abbreviations

AD: Alzheimer’s disease
Aβ: amyloid-beta
FDA: US Food and Drug Administration
AβPP: Aβ protein precursor
EM: electron microscope
Cryo-EM: cryogenic electron microscope
TEM: transmission electron microscope
CAA: cerebral amyloid angiopathy
ARIA-E: amyloid-related imaging abnormalities with edema
PET: positron emission tomography

## Acknowledgements

F.S.P.R. and G.F.S. acknowledge with gratitude the electron microscopy training, imaging and access time provided by the life science electron microscopy facility of the Ernst Ruska-Centre at Forschungszentrum Jülich. The brain tissue from the tg-APP_ArcSwe_, ARTE10, tg-APP_Swe_, APP23, APP/PS1 and tg-SwDI mouse models was generously provided by Dr. Sarah Schemmert, Prof. Dr. Antje Willuweit, Dr. Lili Donner, Prof. Dr. Margitta Elvers, Prof. Dr. Lars N. G. Nilsson, Prof. Dr. Stina Syvänen, Prof. Dr. Dag Sehlin, and Prof. Dr. Martin Ingelsson, who also bred and kept the mice under optimal conditions until euthanized for research purposes. The ARTE10 mouse model was a gift from Taconic Biosciences. D.W. was supported by ‘Portfolio Drug Research’ of the ‘Impuls und Vernetzungs-Fonds der Helmholtzgemeinschaft.’

## Author Contributions

F.S.P.R. conducted the *ex-vivo* Aβ fibril preparations, their lecanemab immunogold-labelling and negative-stain EM of the Aβ fibrils. L.G., D.W. and G.F.S. supervised the project. F.S.P.R. and L.G. wrote the original manuscript draft, F.S.P.R., L.G., G.F.S. and D.W. edited and reviewed the final manuscript.

